# ECM-Focused Proteomic Analysis of Ear Punch Regeneration in *Acomys Cahirinus*

**DOI:** 10.1101/2023.10.11.561940

**Authors:** Maxwell C. McCabe, Daryl M. Okamura, Christopher B. Erickson, Blair W. Perry, Chris M. Brewer, Elizabeth D. Nguyen, Anthony J. Saviola, Mark W. Majesky, Kirk C. Hansen

## Abstract

In mammals, significant injury is generally followed by the formation of a fibrotic scar which provides structural integrity but fails to functionally restore damaged tissue. Spiny mice of the genus *Acomys* represent the first example of full skin autotomy in mammals. *Acomys cahirinus* has evolved extremely weak skin as a strategy to avoid predation and is able to repeatedly regenerate healthy tissue without scar after severe skin injury or full-thickness ear punches. Extracellular matrix (ECM) composition is a critical regulator of wound repair and scar formation and previous studies have suggested that alterations in its expression may be responsible for the differences in regenerative capacity observed between *Mus musculus* and *A. cahirinus*, yet analysis of this critical tissue component has been limited in previous studies by its insolubility and resistance to extraction. Here, we utilize a 2-step ECM-optimized extraction to perform proteomic analysis of tissue composition during wound repair after full-thickness ear punches in *A. cahirinus* and *M. musculus* from weeks 1 to 4 post-injury. We observe changes in a wide range of ECM proteins which have been previously implicated in wound regeneration and scar formation, including collagens, coagulation and provisional matrix proteins, and matricryptic signaling peptides. We additionally report differences in crosslinking enzyme activity and ECM protein solubility between *Mus* and *Acomys.* Furthermore, we observed rapid and sustained increases in CD206, a marker of pro-regenerative M2 macrophages, in *Acomys,* whereas little or no increase in CD206 was detected in *Mus.* Together, these findings contribute to a comprehensive understanding of tissue cues which drive the regenerative capacity of *Acomys* and identify a number of potential targets for future pro-regenerative therapies.

## Introduction

In mammals, tissue damage repair is mediated through a series of biochemical events including inflammatory response, granulation tissue formation, and extracellular matrix (ECM) remodeling^1^. During healing, ECM remodeling typically results in the replacement of healthy tissue with fibrotic scar tissue^2^ which is characterized by excessive deposition of irregularly organized structural ECM proteins, primarily fibrillar collagen^3^. Alternatively, reduced ECM remodeling can lead to altered matrix-regulated autophagy and result in a chronic wound or ulcer^4^. Additionally, ECM composition has been shown to regulate a number of key wound healing processes, including inflammation^5,6^, fibroblast activity^7^, immune cell polarization^8^, and tissue-specific cell differentiation^9^. Therefore, while the tuning of ECM production is critical to the regeneration of healthy tissue, the general inability of mammals to restore healthy tissue after significant injury has driven the increasing prominence of regenerative medicine research, which aims to restore healthy tissue after damage occurs.

The spiny mouse (genus: *Acomys*) has evolved a highly brittle skin that tears easily in response to low applied tension with no fracture plane^10^, unlike geckos and skinks which have a preexisting fracture plane. Tearing is followed by rapid wound contraction (accounting for 95% of wound closure) which results in regeneration of fully functional skin over the wound area. The skin’s low tensile strength and high regenerative capacity appear to have evolved as a mechanisms to evade predators in the wild^10^. Because the mechanical strength of tissue is dictated in large part by its ECM protein composition^11^, it is likely that there are large-scale differences in matrix composition between the skin of *Acomys* and that of *Mus musculus,* which has much stronger skin and lacks the same regenerative capacity. While other examples of regeneration in mammals have been identified (e.g. liver regeneration^12^, embryonic skin^13^), *Acomys* represents the first demonstration of full skin autotomy in mammals. Skin regeneration was first documented in *A. kempi and A. percivali* by Seifert et al. in 2012^10^, and was followed by studies involving full-thickness 4mm ear punches, first in *A. kempi* and *A. percivali*^10^ then later replicated in *A. cahirinus*^14,15^.

Ear punch tissue contains skin, hair follicles, adipose cells, muscle, and cartilage, making it a good model of complex tissue regeneration. In the first analysis of ear punch regeneration, *Acomys* was found to rapidly close these ear punches, regenerating all tissue types with the exception of muscle^10^. However, regeneration of skeletal muscle by *Acomys* in both a later ear punch model^15^ and other contexts has been subsequently documented in other studies^16,17^. In contrast, it has been shown that *Mus musculus* (C57BL/6) heal their wounds through fibrotic scarring with little to no evidence of tissue regeneration. While *M. musculus* ear punches showed no significant closure and appeared fully healed by day 14 post-injury, by day 21 *Acomys* injuries displayed a newly generated ring of translucent tissue which continued to grow inward until 100% of ear punches were fully closed by day 56^15^. *Acomys* regenerative capacity has also been demonstrated in a wide variety of other tissues, including kidney^18^, skeletal muscle^16,17^, cardiac tissue^19–21^, and spinal cord^22,23^. In subsequent regenerative studies, *A. cahirinus* has become the most commonly used experimental species due to its greater accessibility as a model organism. Outside of its regenerative ability, *A. cahirinus* was already in limited use as a laboratory organism due to its other unique biological similarities to humans including propensity for diabetes^24^, menstruation^25^, production of steroids in the adrenal glands^26^, and nephrogenesis before birth^27^.

Previous proteomic studies investigating ear punch regeneration have been conducted in *Acomys*, however, numerous experimental limitations allow for significant improvements. First, earlier studies interpreted *Acomys* proteomic data using the *M. musculus* proteome database which limits the amount of *Acomys* data that can be identified and subsequently the comparisons between *Mus* and *Acomys* that can be performed. Second, previous data suggests large-scale differences in ECM composition between *Mus* and *Acomys,* which may be partly responsible for *Acomys* regenerative capacity^10,14^. However, due to difficulties in solubilizing, and therefore characterizing, the ECM via traditional proteomic methods^28^, a comprehensive characterization of the ECM in *Acomys* is lacking. Consequently, a deeper, more thorough ECM-focused proteomic approach can provide novel insights into the regenerative properties of *Acomys*. Third, previous proteomic analyses of ear punch regeneration involved samples collected only up to day 14 post-injury^14^. Other studies have shown that >85% of wound closure and new tissue growth occurs after day 14 in *Acomys*^15^, indicating that continued sampling after day 14 is essential for capturing the proteome of newly synthesized tissue in *Acomys.* Here, we performed ECM-optimized 2-step extraction for proteomic analysis via LC-MS/MS on the healing edge of full-thickness 4 mm ear punch wounds from *Mus* and *Acomys* sampled weekly from 0 to 4 weeks post-injury. Additionally, we utilize previously published RNA-sequencing data^29^ to generate an annotated *Acomys* protein sequence database for interpretation of the acquired of the mass spectrometric data. The results here provide an overall more comprehensive and more accurate characterization of the ECM allowing for a better understanding of the regenerative capacity of *Acomys*.

## Materials and Methods

### *Acomys cahirinus* Protein Sequence Database Generation

An *A. cahirinus* specific protein database was constructed from a previously published transcriptome^29^ (Tr2aacds_v2.fasta). The TransDecoder.LongOrfs tool in Transdecoder v5.5.0 (github.com/TransDecoder/TransDecoder) was used to extract open reading frames from the transcriptome and translate to putative proteins. The resulting protein sequences were searched against the Uniprot database^30^ (release 2021_03) using BLASTp v2.12.0^31^ with flags - max_target_seqs 1, -outfmt 6, and -evalue 1e-5. TransDecoder.Predict was then run using results from the BLASTp search to filter out spurious proteins with no homology to known proteins. The resulting putative protein set was filtered to retain only complete proteins (those including both a start and stop codon). The rmdup tool in seqkit v2.0.0^32^ was used to identify instances where multiple ORFs were predicted to produce identical proteins, for which a single ORF was retained.

To annotate putative proteins, we used a reciprocal best BLAST (RBB) followed by one-way best BLAST (OBB) approach with the *Mus musculus* protein database (version GRCm39; NCBI: GCA_000001635.9) as the subject database. In instances where a *Mus* protein was assigned an RBB hit in our protein set, all other OBB hits to that *Mus* protein were removed. In instances where multiple proteins had OBB, but not RBB, hits to the same *Mus* protein, the protein with the highest bitscore was retained as the best match and all others removed. This approach resulted in a database of 18,682 putative *Acomys* proteins with homology to *Mus* proteins.

### Sample Acquisition

The Seattle Children’s Research Institute’s (SCRI) Institutional Animal Care and Use Committee (IACUC) approved all animal procedures. Adult male and female *Mus musculus* (The Jackson Laboratory, CD1(ICR)) and *Acomys cahirinus* (colony at SCRI) were maintained within the Seattle Children’s Research Institute’s onsite vivarium. CD1 mice were housed in a pathogen-free room maintained on 12:12 (Light:Dark) lighting schedule, *Acomys* were housed in a separate room maintained on 14:10 (Light:Dark) schedule, and all animals received food and water ad libitum. *Acomys* and *Mus* adult males and females were anaesthetized with 4% (v/v) vaporized isoflurane (Henry Schein Animal Health, Dublin, OH), and a 2-mm thumb punch (Kent Scientific) was used to generate punches across the medial ear pinna in the right and left ears. Punches were then collected from the healing edge of the wound at weeks 1 to 4 post-injury, collecting at only a single post-injury time point for each original injury site. All samples were immediately flash frozen in liquid nitrogen after collection for storage.

### Sample Preparation

Dissected tissue samples were received frozen and lyophilized prior to analysis. Lyophilized samples were cut into <1 mm^2^ pieces using a razor blade and weighed before extraction. Samples were processed using a 2-step extraction beginning with 3 successive washes with decellularization buffer (50mM Tris-HCl (pH 7.4), .25% CHAPS, 25mM EDTA, 3M NaCl). Approximately 100 mg of 3mm glass beads were used to mechanically agitate samples in a Bullet Blender (NextAdvance) prior to all extraction steps. For decellularization, samples were vortexed at 4⁰C for 20 minutes after agitation, centrifuged at 18,000xg for 20 minutes, and the supernatant was collected. Successive decellularization washes were pooled by sample for analysis. After decellularization, sample pellets were subjected to chemical digestion with 1M hydroxylamine hydrochloride (HA) in 4.2M Gnd-HCl, 0.2M K2CO3, pH 9.0 at 42⁰C for 4 hours with constant agitation at 120 RPM on an orbital shaker (Labnet Orbit M60). After digestion, samples were centrifuged at 18,000xg for 20 minutes and the supernatant was collected. This 2-step extraction produced separate cellular and ECM fractions for each sample. Extracted fractions were precipitated using ice-cold 80% acetone (Fisher Scientific #A929-1), washed 2x with ice-cold 100% acetone, and brought up in 8M urea prior to analysis. Protein concentration of each fraction was measured using A660 Protein Assay (Pierce™). Proteolytic digestion of sample extracts was carried out according to the FASP protocol^33^ using 10 kDa molecular weight cutoff filters (Sartorius Vivacon 500 #VN01H02) and loading 10 ug of protein resulting from each fraction. Samples were prepared by reducing, alkylating, and digesting with trypsin (1:100) at 37°C for 14 Hrs. Peptides were recovered from the filter using successive washes with 0.2% formic acid (FA). Samples were desalted using Pierce^TM^ C18 Spin Tips (Thermo Scientific #84850) according to the manufacturer’s protocol. Final volume was adjusted to inject 2 µg of protein.

### LC-MS/MS Analysis

Digested peptides (2 µg) were loaded onto individual Evotips following the manufacturers protocol and separated on an Evosep One chromatography system (Evosep, Odense, Denmark) using the default 30 samples per day LC method over a Pepsep column (150 um inter diameter, 15 cm) packed with ReproSil C18 1.9 um, 120A resin. The system was coupled to the timsTOF Pro mass spectrometer (Bruker Daltonics, Bremen, Germany) via the nano-electrospray ion source (Captive Spray, Bruker Daltonics). The mass spectrometer was operated in PASEF mode. The ramp time was set to 100 ms and 10 PASEF MS/MS scans per topN acquisition cycle were acquired. MS and MS/MS spectra were recorded from m/z 100 to 1700. The ion mobility was scanned from 0.7 to 1.50 Vs/cm2. Precursors for data-dependent acquisition were isolated within ± 1 Th and fragmented with an ion mobility-dependent collision energy, which was linearly increased from 20 to 59 eV in positive mode. Low-abundance precursor ions with an intensity above a threshold of 500 counts but below a target value of 20000 counts were repeatedly scheduled and otherwise dynamically excluded for 0.4 min.

### Data Analysis

Raw data was searched using MSFragger (version 3.5) through the FragPipe platform (version 18.0). Precursor tolerance was set to ±15 ppm and fragment tolerance was set to ±0.08 Da, allowing for 2 missed cleavages. *M. musculus* data was searched against SwissProt (17,029 sequences) restricted to *M. musculus*, while *A. cahirinus* data was searched against the custom sequence database described above (18,682 sequences). All databases had common contaminants added using version 1.1 of the CRAPome^34^. Trypsin semi-specific cleavage was used in all searches. Fixed modifications were set as carbamidomethyl (C). Variable modifications were set as oxidation (M), oxidation (P) (hydroxyproline), Gln->pyro-Glu (N-term Q), deamidated (NQ), and acetyl (protein N-term). Results were filtered to 1% false discovery rate (FDR) at the peptide and protein level using Philosopher (version 4.4.0) employed through FragPipe. Label-free quantification (LFQ) was performed using the IonQuant (version 1.8.0), including unique and razor peptides. Cell and ECM fractions for each organism were searched independently and data was normalized to total LFQ intensity by fraction, so that total signal in each fraction was equal across all samples before fractions were aggregated for analysis. Reported core matrisome (ECM proteins) and matrisome associated proteins were annotated with Matrisome DB^35^.

## Results and Discussion

To improve identification of *Acomys* peptides non-homologous to *Mus* sequences, we first generated a protein sequence database using previously published RNA-sequencing (RNA-seq) data^29^ (Figure 1). In brief, open reading frames were extracted and translated from RNA-seq data before performing a homology search against UniProt with unrestricted taxonomy. Multiple filtering steps were performed to remove aberrant transcripts before annotating protein sequence based on their homology to *M. musculus* protein entries. Finally, the database was filtered to remove all but the most complete sequence for each protein, resulting in 18,682 protein sequences (Figure 1).

**Figure 1.**
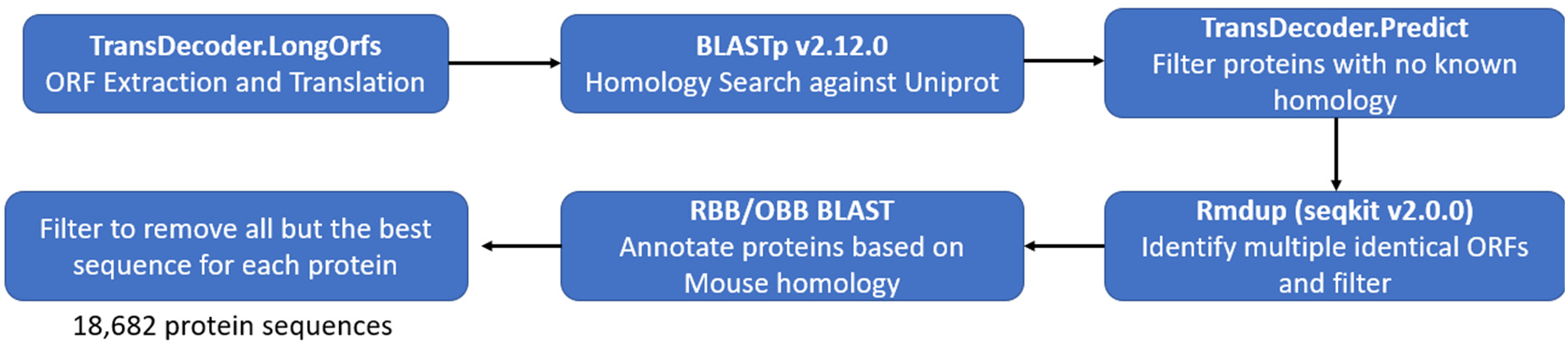
Workflow for annotated protein database generation from RNA-seq data.

To assess the quality of our newly generated *Acomys* protein database, a comparison of *Acomys* MS data searched against both the *M. musculus* SwissProt database (17,268 sequences), and in-house *A. cahirinus* (18,682 sequences) database was performed (Table 1). The *A. cahirinus* database resulted in a 47% increase in total peptides and nearly a 2-fold increase in ECM peptides, demonstrating significant improvement in peptide IDs across the proteome. Additionally, *Acomys* data searched against the in-house DB provides an additional 1,088 peptide sequences and 983 ECM peptide sequences compared to *Mus* data searched against the *Mus* database. However, while the *Acomys-*specific DB results in 142 additional protein identifications for *Acomys* data compared to the *Mus* DB, the best-performing *Acomys* data analysis results in only 67% of total and 80% of ECM *Mus* protein identifications (Table 1).

**Table 1.** Comparison of *Acomys* and *Mus* database performance for *Acomys* data analysis.

|  | *Acomys* vs *Mus* DB<br>(previous approach) | *Acomys* vs *Acomys* DB<br>(current approach) | *Mus* vs *Mus* DB<br>(gold standard) |
| --- | --- | --- | --- |
| Total Peptides Identified | 53566 | 78951 | 77863 |
| ECM Peptides Identified | 5850 | 11495 | 10512 |
| Total Proteins Identified | 4652 | 4794 | 6154 |
| ECM Proteins Identified | 252 | 264 | 330 |

This study utilized the model of ear punch regeneration employed on *Acomys* and *Mus* as described by Brewer et al.^36^ coupled with an ECM-focused extraction method to provide a comparison of protein alterations during wound healing between *Acomys* and *Mus* (Figure 2). In brief, 2-mm punches were taken from the central region of *A. cahirinus* and *M. musculus* ears (n = 24). At weeks 1 to 4 post injury, the healing edge of the wound was excised, collecting the healing edge at only one time point from each mouse (n=4-6 per time point). Samples were then subjected to a 2-step ECM-focused protein extraction method, consisting of decellularization using CHAPS and 3M NaCl followed by ECM extraction using hydroxylamine-HCl (HA) and guanidine-HCl (Gnd-HCl). Extracted fractions were analyzed via LC-MS/MS, resulting in 4,794 protein IDs for *Acomys* and 6,154 protein IDs for *Mus*.

**Figure 2.**
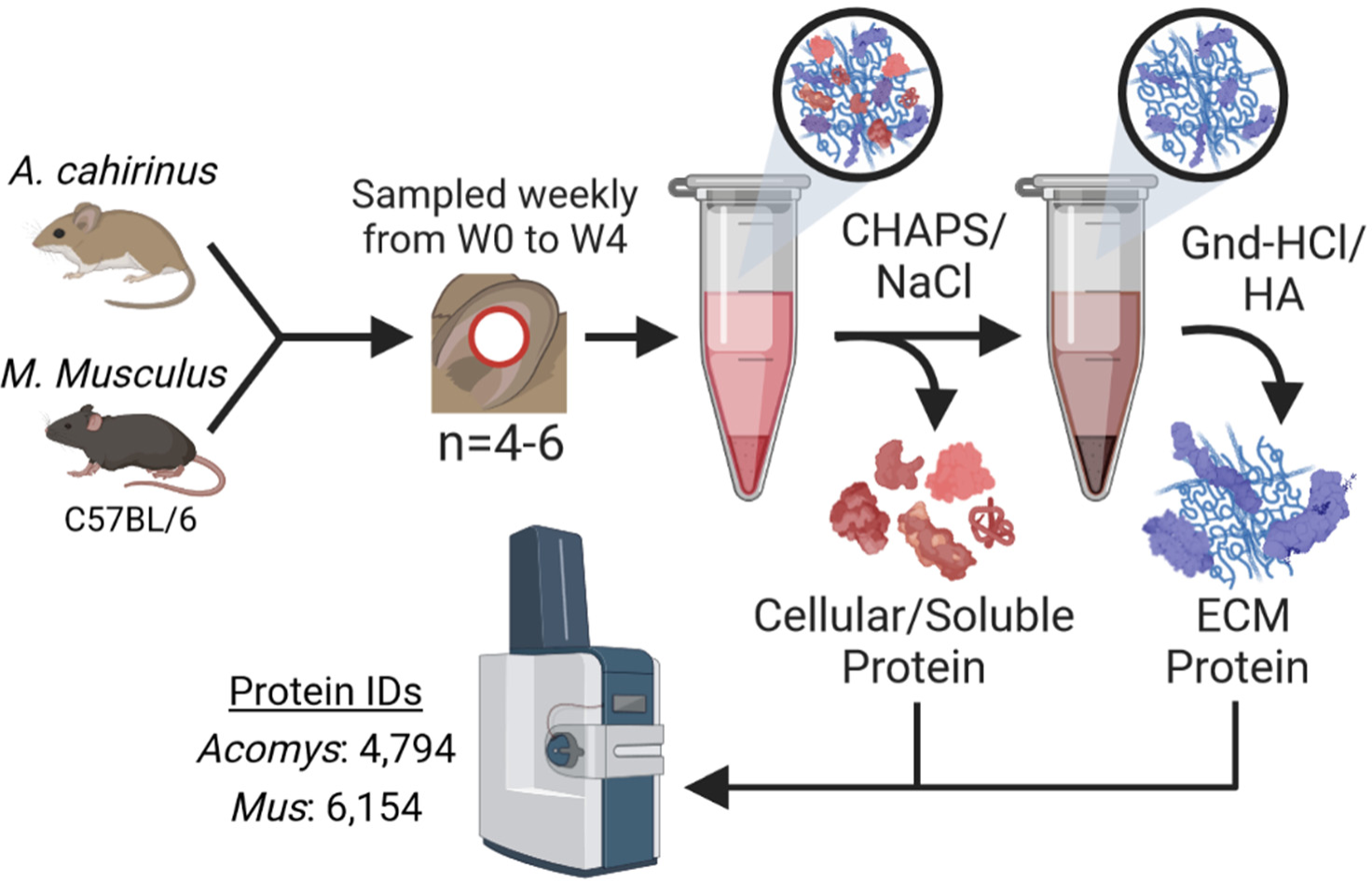
Workflow for proteomic analysis of ear punch regeneration in *A. cahirinus* and *M. musculus* (C57BL/6). Full-thickness 4 mm ear punches were performed on *A. cahirinus* and *M. musculus* (n=4-6 per time point) and the excised tissue was collected (W0). Tissue was then collected from the healing edge of the ear punch wounds weekly from W1 to W4. Samples were subjected to a 2-step ECM-optimized extraction, digested, and analyzed using a timsTOF Pro MS system.

To establish baseline differences between the proteomes of *Mus* and *Acomys*, comparisons were performed between initial ear punch samples containing uninjured tissue (Figure 3). Significant differences are observed in a wide range of cellular proteins, including cytoskeletal components (actins, tubulins, and cytoskeletal keratins) and metabolic enzymes (Figure 3A). While not the primary focus of this study, cellular protein alterations between uninjured *Mus* and *Acomys* tissue indicate baseline differences in metabolic capacity and cellular structure between the two species.

**Figure 3.**
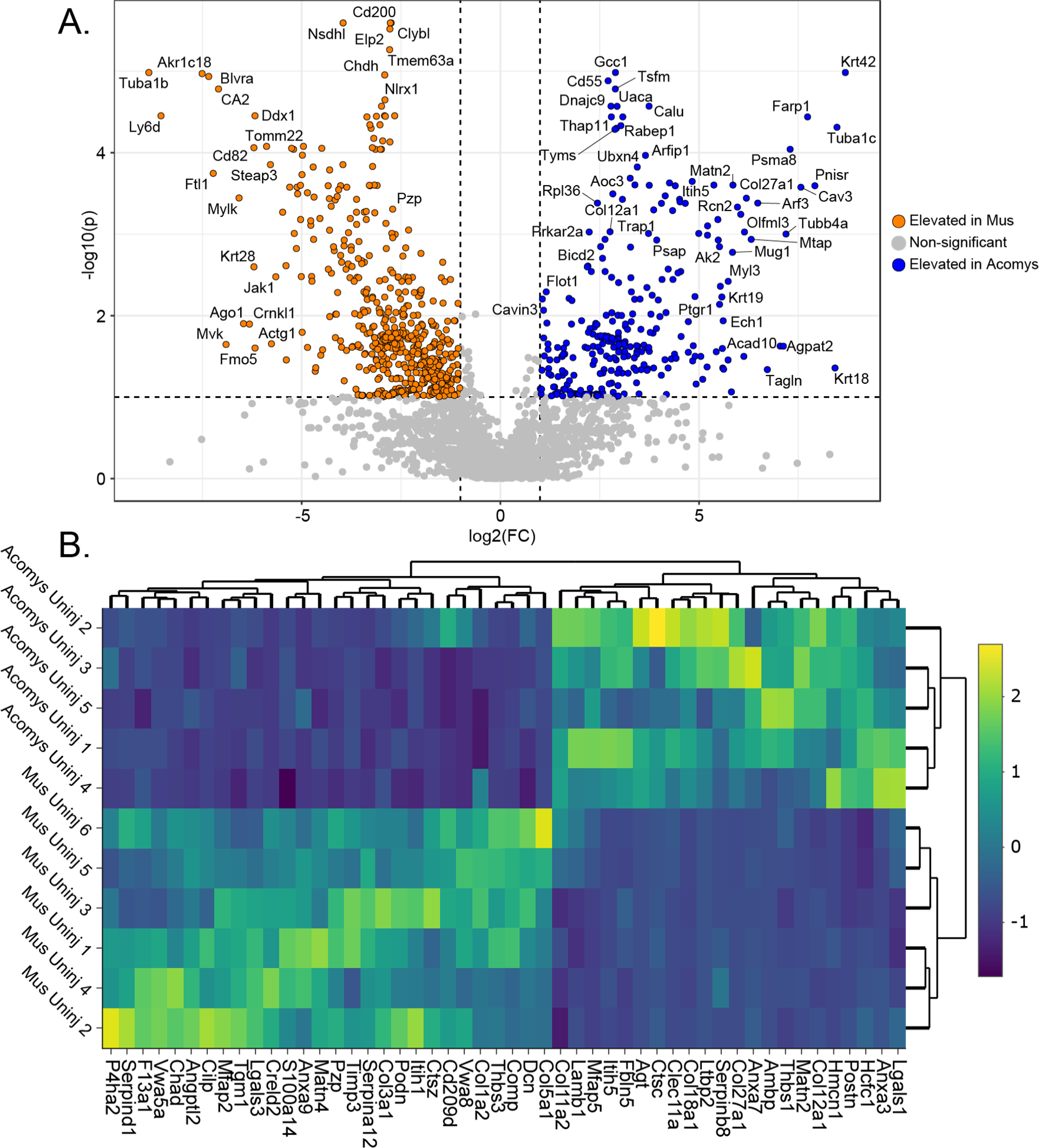
Baseline comparison of uninjured *Mus* and *Acomys.* A) Volcano plots showing all proteins significantly elevated in uninjured *Mus* (left, orange) and *Acomys* (right, blue). B) Heatmap displaying top 50 distinguishing ECM proteins by t-test between uninjured *Mus* and *Acomys*.

Large-scale differences in ECM protein composition are also observed between uninjured *Mus* and *Acomys* (Figure 3B). We observe elevated expression of fibrillar collagens COL1A2 and COL5A1 in *Mus,* alongside collagen-supporting proteoglycan decorin (DCN), indicating a more robust fibrillar collagen network in *Mus* which likely contributes to the greater tensile strength of *Mus* tissue. Of the collagens most upregulated in *Acomys* compared to *Mus,* including COL4A5, COL5A3, COL6A3, COL9A1, COL9A3, COL11A2, COL12A1, COL18A1, and COL27A1, only two are fibrillar (COL5A3 and COL27A1), with the rest represented by FACIT, network, and other collagens. The increased abundance of many collagen subtypes which are typically considered “minor” in most organisms^37^. Many differences in non-collagen matrix proteins are also detected, including elevated galectin-3 (LGALS3) and chondroadherin (CHAD) in *Mus* and elevated matrillin-2 (MATN2), and laminin chains in *Acomys* (Figure 3B). Many differentially regulated ECM proteins between uninjured *Acomys* and *Mus* have been implicated in either pro-regenerative or pro-inflammatory processes during wound repair (Table 2).

**Table 2.** Selected differentially expressed proteins enriched in *Acomys* (top) and *Mus* (bottom) with identified roles in wound healing and regeneration.

| Gene | *Acomys* Expression | *Mus* Expression | Wound Healing Role | KO Effect |
| --- | --- | --- | --- | --- |
| POSTN | High | Low, moderate induction after injury | Regulates COL fibrillogenesis and skin viscoelasticity<sup>46</sup>/Associated with myofibroblast transition<sup>47</sup> | Impaired wound closure<sup>47</sup>, reepithelialization<sup>48</sup>, and COL crosslinking<sup>46</sup> |
| FBLN2 | Low in uninjured, strong induction after injury | Low, weak induction after injury | Maintains basement membrane integrity<sup>49,50</sup>, participates in formation of provisional FN1 matrix<sup>51</sup> | Skin blistering, reduced basement membrane integrity<sup>50</sup> |
| MATN2 | High, low at week 1 with later recovery | Very low in uninjured, moderate induction after injury | Multiadhesive adaptor protein essential for the regeneration of muscle, nerve, skin, and other tissues<sup>52</sup> | Myoblast cell cycle arrest, delayed muscle regeneration<sup>53</sup> |
| AEBP1 | High | Low in uninjured, moderate induction at week 2 | Regulate fibroblast proliferation, COL production and MSC differentiation<sup>54</sup> | Delayed wound repair correlating with defects in fibroblast proliferation<sup>54</sup> |
| THBS1 | High, increased at weeks 2 and 3 | Low, slight induction after injury | Activates TGF-β, regulates angiogenesis<sup>55</sup> and COL assembly, mediates ER stress response<sup>56</sup> | Delayed repair, loosely compacted and disorganized granulation tissue<sup>55</sup> |
| HSPG2 | High in uninjured, lower at week 1 with some recovery | Moderate in uninjured, lower after injury | Orchestrates morphogen signaling<sup>42</sup>, stabilizes ECM, promotes angiogenesis<sup>43</sup> | Delayed wound healing in deficient mice<sup>57</sup> |
| C1QC | Low in uninjured, strong induction after injury | Low in uninjured, moderate induction after injury | Supports new vessel formation, promotes wound healing as DDR2 ligand<sup>58</sup> | Defective vessel formation, lower vessel density, impaired wound healing<sup>59</sup> |
| PODN | Moderate | High, increased at week 4 | Negative regulator of cell migration and proliferation<sup>60</sup> | Increased smooth muscle cell proliferation<sup>61</sup> |
| TGM1 | Moderate | High | Crosslinks cornified envelope<sup>38</sup> | Impaired skin barrier function<sup>62</sup> |
| SDC1 | Not detected in uninjured, slight induction after injury | Low in uninjured, strong induction after injury | Modulates integrin activation, upregulated during fibrosis<sup>63</sup> | Delayed wound healing<sup>63</sup> |
| CHAD | Low, weak induction after injury | High in uninjured, Moderate after injury | Binds collagen and inhibits fibrillogenesis in vitro, reduces chondrocyte differentiation<sup>64</sup> | Increased chondrocyte differentiation and collagen II deposition<sup>64</sup> |
| ADAM9 | Very low | Low in uninjured, strong induction after injury | Causes collagen XVII ectodomain shedding<sup>65</sup> | Accelerated wound repair and keratinocyte migration<sup>66</sup> |
| COL14A1 | Low | Low in uninjured, strong induction at week 2 | Regulation of collagen fibrillogenesis<sup>67</sup> | Reduced tensile strength of tissue<sup>67</sup> |

Analysis of ECM protein expression during the course of wound healing in *Acomys* and *Mus* reveals functionally significant clusters of differentially expressed proteins between organisms (Figure 4). Cluster 1 reveals proteins which are expressed at moderate to low intensity in *Acomys* and high intensity in *Mus,* with many proteins stably expressed and some decreasing in abundance over the course of wound healing. These proteins include fibrillar collagens COL1A2, COL3A1, and COL5A1, all of which have been associated with and are overexpressed in fibrosis and scar formation^2^. We also observe cornified envelope protein S100A14 as well as transglutaminase 1 (TGM1), the principal enzyme responsible for crosslinking of the cornified envelope^38^, in this cluster.

**Figure 4.**
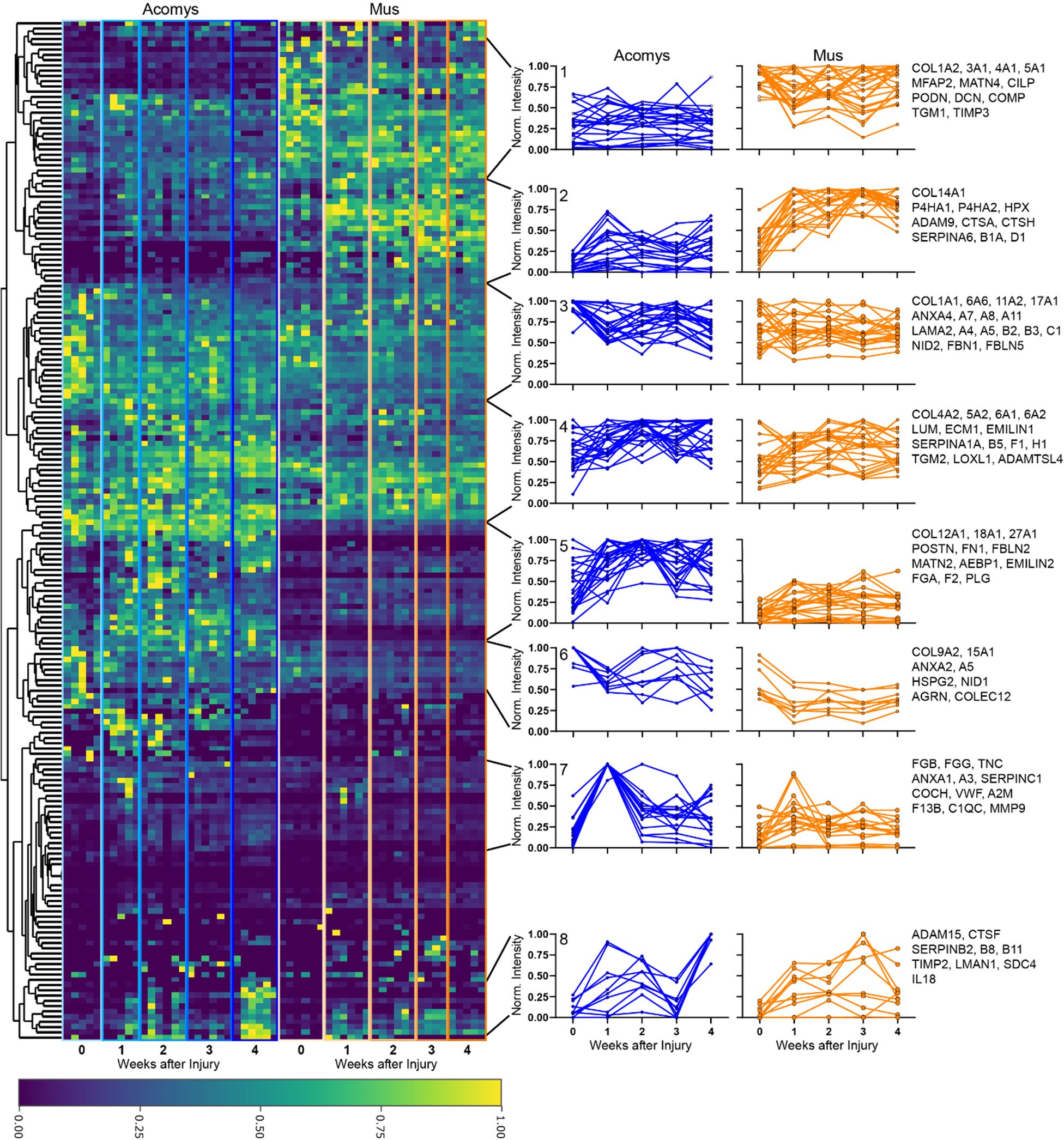
Expression profiles of matrisome proteins throughout wound healing reveal functionally significant expression clusters. Heatmap displaying normalized intensity for all core matrisome proteins across *Acomys* (blue) and *Mus* (orange) samples from uninjured (0 weeks after injury) to week 4 (left). Profile plots displaying average intensity (normalized to maximum intensity for each protein across all samples) from uninjured to week 4 post-injury in *Acomys* (blue) and *Mus* (orange) for all identified proteins in the specified cluster (right). Cluster numbers are indicated in the upper-left corner of *Acomys* profile plots.

Cluster 2 shows proteins elevated in *Acomys* compared to *Mus*. However, in this cluster we observe induction of many proteins after injury in both species, with intensity remining high throughout the course of wound healing in *Mus* but not in *Acomys.* Proteins assigned to this cluster include a variety of proteases involved in ECM remodeling (cathepsin A (CTSA), cathepsin H (CTSH), disintegrin and metalloproteinase domain-containing protein 9 (ADAM9)) as well as ECM protease inhibitors (Inter-alpha-trypsin inhibitor heavy chain H2 (ITIH2), corticosteroid-binding globulin (SERPINA6), leukocyte elastase inhibitor A (SERPINB1A), heparin cofactor 2 (SERPIND1)). Of note, ADAM9 knockout in mice has been shown to accelerate cutaneous wound healing^39^. Additionally, prolyl hydroxylases A1 and A2 (P4HA1 and P4HA2) are observed in this cluster and are involved in the generation of essential post-translational modifications (PTMs) during collagen synthesis. The persistence of enzymes related to ECM remodeling and synthesis over the course of wound healing in *Mus* but not *Acomys* suggests a longer duration of ECM deposition and remodeling after injury in *Mus.* Clusters 3 and 4 represent proteins which are expressed at high levels in both species throughout the course of wound repair. These proteins include a variety of collagen and laminin chains, as well as annexins and ECM crosslinking enzymes (transglutaminase 2 (TGM2), lysyl oxidase homolog 1 (LOXL1)) (Figure 4).

Cluster 5 displays proteins which are significantly elevated in *Acomys* compared to *Mus*, with generally larger induction after injury in *Acomys* (Figure 4). This cluster contains a number of proteins which have been associated with a pro-regenerative phenotype or whose absence has been implicated in impaired wound healing in previous studies, including periostin (POSTN), fibulin-2 (FBLN2), matrillin-2 (MATN2), adipocyte enhancer-binding protein 1 (AEBP1), and thrombospondin-1 (THBS1) (Table 2). We additionally observe a number of proteins related to coagulation (fibrinogen alpha chain (FGA), plasminogen (PLG), prothrombin (F2)), early wound healing, and provisional matrix deposition (fibronectin (FN1)) in this cluster, suggesting a more robust early wound healing response in *Acomys* compared to *Mus*. FN1 plays a key role in the formation of early provisional matrix, and FBN2 has previously been shown to colocalize and coexpress with FN1 during early stage of skin regeneration, forming a critical component of the provisional FN1 matrix^40^.

Cluster 6 contains proteins which are detected at lower levels at week 4 post-injury than in uninjured tissue in both species, with generally higher expression in *Acomys* throughout the course of wound repair. These proteins include basement membrane components nidogen-1 (NID1) and perlecan (HSPG2), both of which are essential for the construction and maintenance of the basement membrane. NID1 deficiency during skin wound healing was found to induce hyperproliferation and differentiation delays as well as aberrant laminin deposition and distribution, demonstrating a critical role of NID1 during tissue repair^41^. HSPG2, as well as being a critical structural component of the basement membrane, regulates the binding of morphogens and mitogens to cells^42^, altering signaling pathways and cell behavior, assists with collagen fibrillogenesis^42^, and promotes angiogenesis^43^. HSPG2 is highly conserved across multicellular organisms and has been found to be critical for wound repair in the *Nematostella vectensis* model of regeneration^44^.

In cluster 7, we observe proteins which are present at low levels in uninjured samples in both species. In *Acomys*, all proteins contained in this cluster show significant upregulation after injury coupled with recovery toward uninjured levels after week 1 post-injury, while in *Mus* upregulation is observed in only a subset of these proteins. All proteins in this cluster are identified at the highest intensity in *Acomys* week 1 post-injury samples except for tenascin-C (TNC), which is identified at the greatest intensity in *Acomys* week 2 post-injury. Proteins in this cluster are highly enriched in components of the coagulation cascade, including fibrinogen beta chain (FGB), fibrinogen gamma chain (FGG), coagulation factor XIII B chain (F13B), cochlin (COCH), and von Willebrand factor (VWF). The increased expression of a wide variety of coagulation factors at week 1 post injury in *Acomys* compared to *Mus* demonstrates a more significant coagulation response in *Acomys* which persists for at least one week. This upregulation of coagulation cascade proteins may contribute to a more robust early wound healing response, representing the formation of a critical initial matrix which stabilizes wound fields, promotes cell migration and proliferation, and ultimately sets the stage for effective regeneration to occur^45^. We additionally observe significant upregulation of TNC in *Acomys* compared to *Mus* which persists after coagulation proteins have dissipated.

Collagens represent the primary structural component of the ECM and play a critical role in both regeneration and the development of fibrosis after injury. Analysis of collagen abundance during wound healing reveals many significant differences in collagen subtype expression between *Acomys* and *Mus* (Figure 5). Scar formation is typically characterized by the deposition of disordered fibrillar collagen^68^. We observe significantly greater abundance of fibrillar collagen in *Mus* compared to *Acomys* in uninjured samples as well as weeks 1 and 2 post injury. However, increased deposition of fibrillar collagen during scar formation is unexpectedly not observed in *Mus* (Figure 5A). This trend does not hold true for all fibrillar collagens, with fibrillar collagens COL3A1 and COL27A1 displaying expression patterns which are distinct from one another as well as from that of all fibrillar collagens combined (Figure 5C, E). While in other models of scarless healing, such as fetal tissue regeneration, COL3A1 is consistently reported to be expressed at higher levels in tissue which performs scarless regeneration than in fibrotic tissue^1,69,70^, we observe elevated COL3A1 in *Mus* compared to *Acomys* (Figure 5C). Like other fibrillar collagens, COL3A1 contributes significantly to tissue tensile strength and knockouts or mutations of the gene in mouse skin cause thin and fragile skin which undergoes increased wounding^71^. The relative lack of COL3A1 in *Acomys* likely contributes to reduced skin strength and suggests that *Acomys* regeneration occurs through mechanisms independent of collagen fibril diameter regulation by COL3A1. Fibrillar collagen COL27A1, on the other hand, is detected at great abundance in *Acomys* at all time points (Figure 5E). COL27A1 is unique from other fibrillar collagens due to its short, interrupted main helical domain^72^ and, due to its relatively recent discovery^73^, little is known about its function in the context of wound healing. Studies have shown that COL27A1 is expressed at high levels in developing cartilage^73^ and its expression is regulated by SRY-Box transcription factor 9, a key regulator of chondrogenesis^74^. Thus, elevated COL27A1 in *Acomys* likely indicates greater representation of this protein in uninjured tissue and improved regeneration of cartilage after injury.

**Figure 5.**
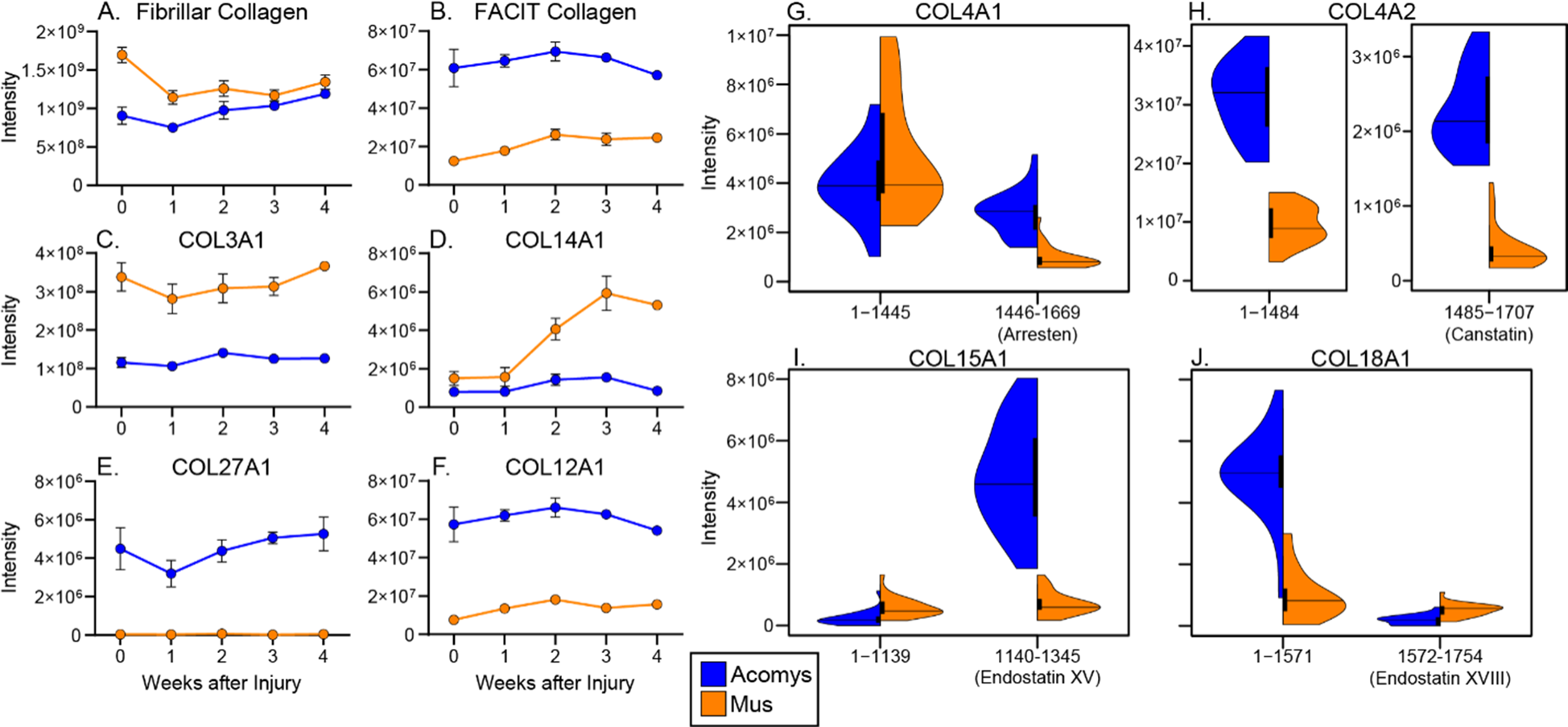
Collagen composition during wound healing varies between *Acomys* and *Mus* and collagen signaling peptides are regulated independently from their parent proteins. Summed intensity for fibrillar collagens (A), FACIT collagens (B), COL3A1 (C), COL14A1 (D), COL27A1 (E), and COL12A1 (F) in *Acomys* (blue) and *Mus* (orange) across the course of wound healing. Uninjured samples are labeled as 0 weeks after injury. G) Intensity for COL4A1 residues 1-1445 (left) and residues 1446-1669 (arresten, right) between *Acomys* and *Mus* across all samples. H) Intensity for COL4A2 residues 1-1484 (left) and residues 1485-1707 (canstatin, right). I) Intensity for COL15A1 residues 1-1139 (left) and residues 1140-1345 (endostatin XV, right). J) Intensity for COL18A1 residues 1-1571 (left) and residues 1572-1754 (endostatin XVIII, right).

Fibrillar-associated collagens with interrupted triple helices (FACIT collagens), which bind the exterior of collagen fibrils to regulate fibrillogenesis and confer various biological functions, display the opposite trend, with strong enrichment in *Acomys* throughout the analyzed time course (Figure 5B). In contrast with this pattern, induction of FACIT collagen COL14A1 is observed after week 1 post-injury in *Mus* but not in *Acomys.* COL14A1 has been shown to contribute to tissue resilience^67^ and is expressed at high levels during porcine fibroproliferative scarring after deep cutaneous injury in pigs^75^, indicating that it may play a role in *Mus* scar formation as well. COL12A1, a FACIT collagen which binds type I collagen fibrils to regulate fiber assembly and adhere other ECM proteins to the fibrillar matrix^76^, is detected at significantly higher abundance in *Acomys* than in *Mus* throughout the course of wound healing (Figure 5F). Knockout of COL12A1 has been shown to delay wound healing through failed sequestration of TGF-β, while overexpression delays healing through increased persistence of M1 macrophages^76^.

The C-terminal non-collagenous domains of type IV, XV, and XVIII collagens can function independently from the structural roles of their parents proteins as potent inhibitors of angiogenesis, motility, and other pro-proliferative cell behaviors^77,78^. COL4A1 and COL4A2 form the primary structural components of the basement membrane and contain non-collagenous signaling domains termed arresten^79^ and canstatin^80^, respectively. Both arresten and canstatin have been shown to inhibit tube formation and proliferation in many types of endothelial cells^77^, reduce both PI3K/Akt activation and vascular endothelial growth factor (VEGF) expression^81^, and induce endothelial cell apoptosis in certain contexts^80,82^. While we observe no significant difference between *Acomys* and *Mus* in abundance of the primary collagenous region of COL4A1 (residues 1-1445), we detect significantly elevated abundance of peptides corresponding to arresten (residues 1446-1669) (Figure 5G). In COL4A2, on the other hand, we observe significant elevation in *Acomys* of both canstatin (residues 1485-1707) and the remainder of the parent protein, with no evident difference in expression pattern between the two regions (Figure 5H).

The α1 chains of type XV and XVIII collagens can also be cleaved to produce molecules termed endostatins which primarily function to inhibit angiogenesis^83,84^. However, key differences have been identified between endostatin-XV (derived from COL15A1) and endostatin-XVIII (derived from COL18A1) which drive the distinct biological functions reported for these highly related proteins^85^. Endostatins XV and XVIII vary significantly in their affinity for common ECM ligands and endostatin-XV lacks the zinc and heparin binding ability which is present in endostatin XVIII and required for inhibition of FGF-2-induced angiogenesis^86^. Additionally, endostatins vary in their localization within tissue structures: endostatin-XVIII is expressed more within elastic fibers and forms a distinct component of portal veins while endostatin XV is a much more prominent component in muscle tissue^86^. We observe greater intensity for the collagenous region of COL15A1 (residues 1-1139) in *Mus*, contrasting with a strong enrichment of endostatin-XV (residues 1140-1345) in *Acomys* (Figure 5I). Endostatin-XVIII displays the opposite trend, showing significant elevation in *Mus* while its parent protein is strongly enriched in *Acomys* (Figure 5J).

While numerous studies have been performed analyzing the effects of endostatin-XVIII in various contexts, including wound healing, much less is known about the independent *in vivo* function of endostatin-XV^85^. Endostatin-XVIII overexpression has been shown to result in delayed wound healing^87^, while deposition of full-length COL18A1 has been demonstrated in the basement membrane during re-epithelialization of mouse skin wounds^88^. The dual role of COL18A1 during skin regeneration is mirrored in our data, where in *Acomys* the parent COL18A1 protein is present at much higher levels than the C-terminal endostatin fragment (Figure 5J). This expression pattern likely allows for COL18A1 to participate in basement membrane formation without inducing the anti-proliferative effects caused by endostatin-XVIII. Further studies are required to elucidate the role of endostatin-XV in *Acomys* regenerative capacity.

Analysis of protein solubility between species over the course of wound healing reveals two significant trends: (1) generally decreased protein solubility in uninjured tissue compared to late wound-healing time points and (2) lower solubility in *Acomys* compared to *Mus,* especially in uninjured tissue. Lysyl oxidase (LOX) is responsible for crosslinking of collagen and elastin fibrils, decreasing their solubility in the process^89,90^. In *Acomys*, LOX significantly increased in abundance at every analyzed time point (Figure 6B), which was coupled with significantly reduced solubility in *Acomys* compared to *Mus* in uninjured (p=0.016), week 1 (p=0.00006), week 2 (p=0.027), and week 3 (p=0.00008) time points (Figure 6C). Collagen crosslinking is usually associated with tissue strength^91,92^; however, we observed higher levels of LOX in *Acomys* where skin tensile strength is very low. We hypothesize that while altered expression of collagen subtypes likely contributes to reduced tensile strength of *Acomys* skin (Figure 5), increased crosslinking may help maintain tissue integrity and architecture during normal physical activity. Additionally, multiple studies have shown that up-regulation of LOX during the early stages of wound healing can result in accelerated wound healing^93,94^. Insufficient levels of LOX during granulation tissue formation can lead to scarring^92^ and LOX has been shown to play an important role in the regulation of angiogenesis^95,96^, an important process for supplying the granulation matrix with nutrients necessary for regeneration^97^.

**Figure 6.**
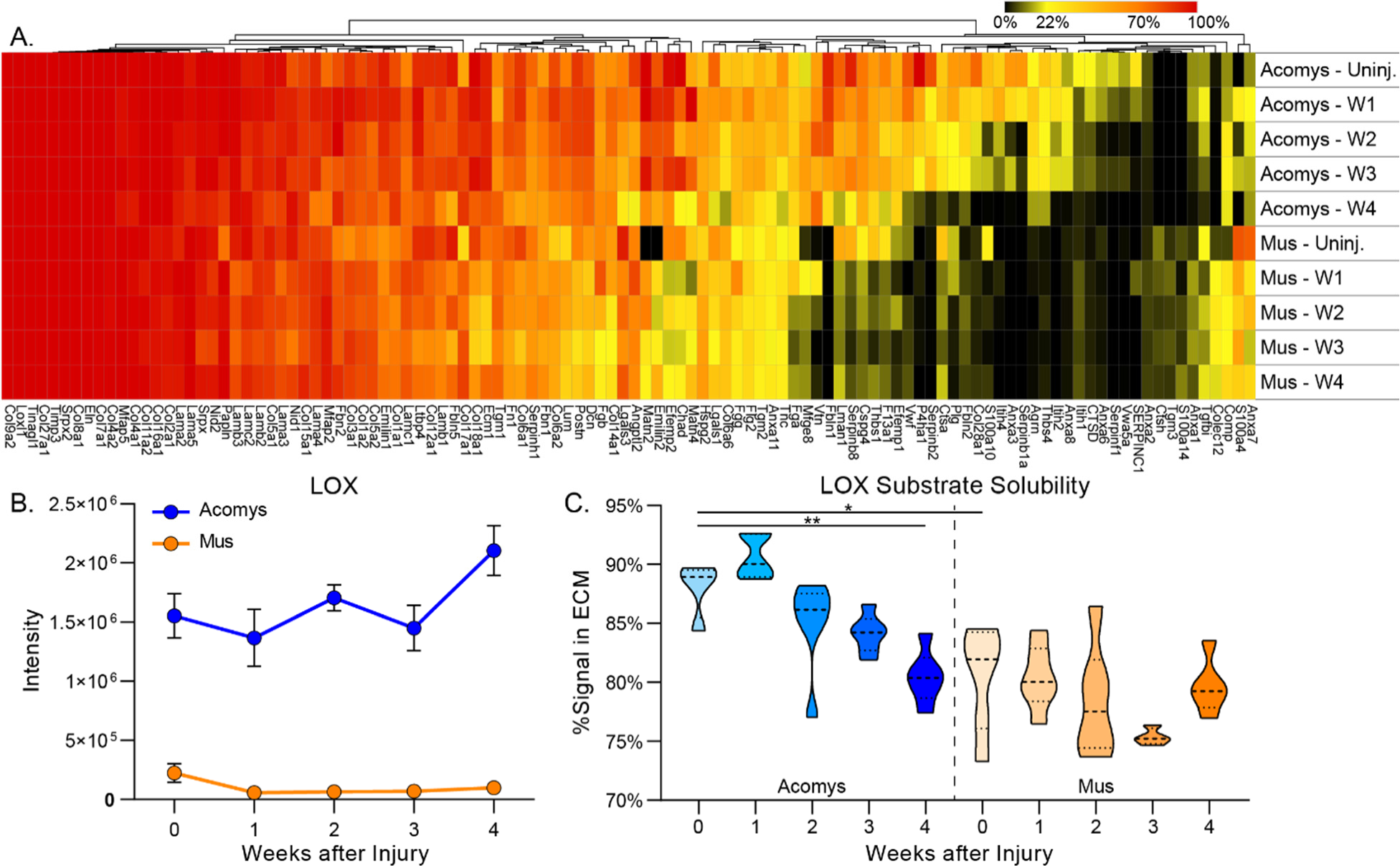
Solubility of LOX substrates and other ECM proteins varies by species and time point. A) Heatmap displaying solubility of ECM proteins measured by the percentage of total signal identified in the ECM fraction for each protein. B) Abundance of LOX in *Mus* and *Acomys* throughout the course of wound healing. C) Average solubility of LOX substrates in *Mus* and *Acomys* over the course of wound healing.

While LOX-family enzyme activity is typically increased during scar formation, both hypertrophic and keloid scar tissue actually display less enzymatic collagen crosslinking than does healthy tissue or mature scar^68^. Similarly, in a model of postburn scarring, LOX was found at low activity in granulation tissue with a sharp increase after 2-3 months, with elevated activity maintained in the tissue for 3-5 years^98^. It is possible that our methods were unable to capture the elevation in LOX activity and expression which occurs at later wound healing time points during scar formation in *Mus*.

In addition to a general trend of decreased solubility at later time points in many ECM proteins (Figure 6A), we observed a significant reduction in LOX substrate solubility in *Acomys* week 4 samples compared to uninjured samples (Figure 6C). The presence of a significant solubility difference between uninjured and week 4 tissue in *Acomys* but not in *Mus* likely demonstrates the increased production of immature, incompletely crosslinked matrix during *Acomys* wound healing. Interestingly, in *Acomys* we detect LOX at the highest abundance in week 4, when protein solubility is highest. This increased LOX abundance after full wound closure may be responsible for crosslinking and regenerating the tensile strength of newly synthesized tissue in the wound area. Previous studies have shown that although 100% of *Acomys* ear punches closed by day 56 post-injury, cartilage architecture remains histologically immature at this point, with morphologically mature cartilage observed at 3 months post-injury^15^. LOX crosslinking activity plays a key role in continued tissue remodeling after wound closure^92^, with the effects of this activity on solubility accumulating over time as crosslinks are formed. It is likely that analysis of later time points would reveal recovery toward uninjured solubility levels as tissue maturation continues.

Analysis of key ECM component and cell marker expression throughout the wound healing trajectories of *Acomys* and *Mus* reveals important insights about how wound repair differs in these two models (Figure 7). Notably, *Acomys* exhibited a more rapid, pronounced, and prolonged increase in Fibrinogen gamma chain (FGG) levels (Figure 7B), suggesting an enhanced clotting response, supported by similar patterns in other clotting proteins (FGA and FGB). As previously mentioned, increased fibrin deposition in *Acomys* may represent the formation of a critical initial matrix which aids in effective tissue regeneration^45^. TNC is a key member of the early provisional matrix formation during wound healing. We found that *Acomys* displayed a more robust induction of TNC after week 1 post-injury compared to *Mus*, where TNC induction was much less pronounced (Figure 7C). This significant difference in TNC expression suggests that *Acomys* may establish a more substantial provisional matrix, which has been shown to play a key role in fetal scarless wound repair^70^ and to promote regeneration in a variety of other contexts^99^. Type 1 collagen is a primary structural component of scar tissue and is generally increased during scar formation. Interestingly, we observed a greater recovery of COL1 levels toward baseline after injury in *Acomys* compared to *Mus* (Figure 7D). This may indicate that *Acomys* possess a heightened ability to remodel and resolve scar tissue, potentially contributing to their regenerative advantage.

**Figure 7.**
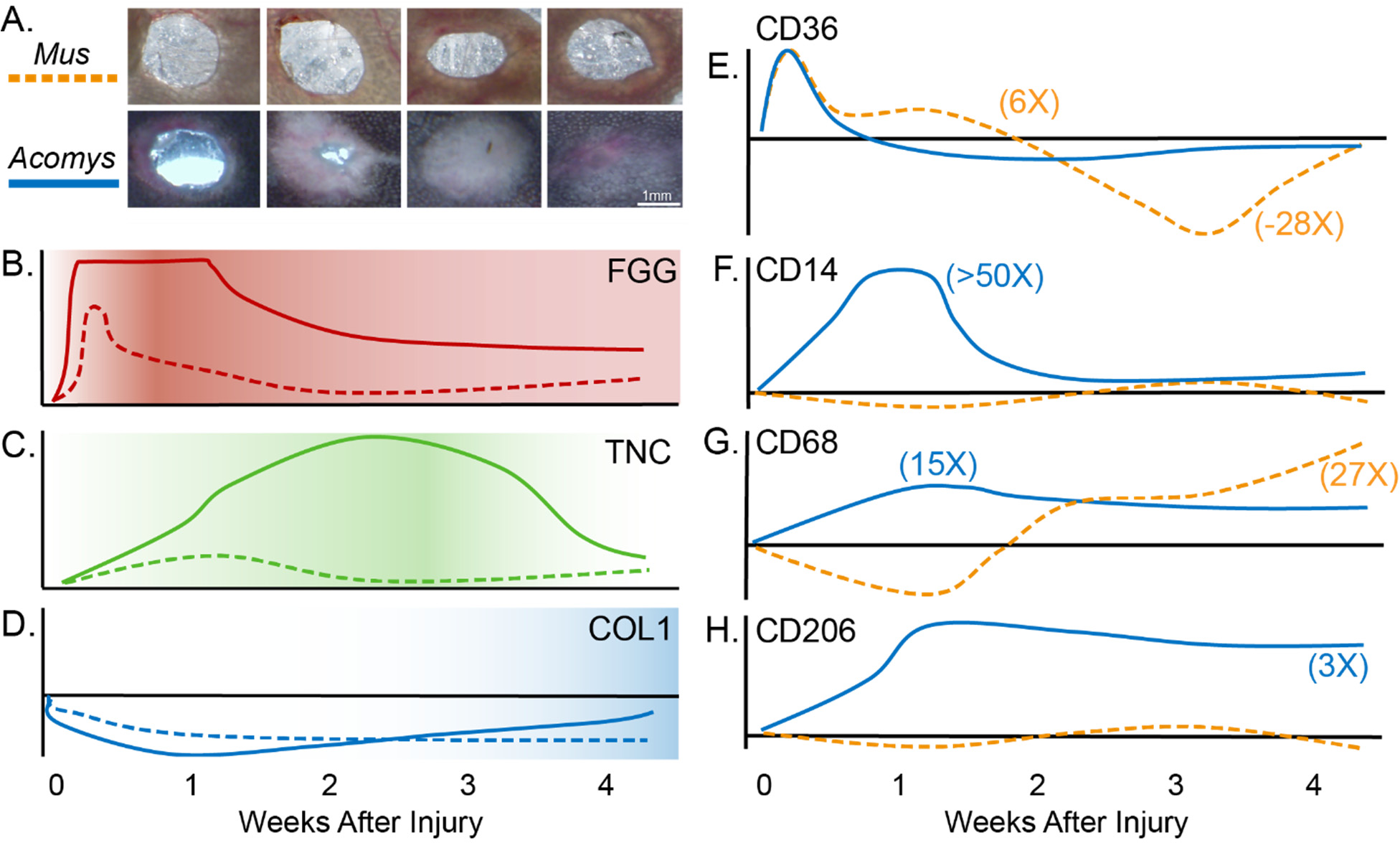
Wound healing trajectory for *Acomys* and *Mus*. A) Images of ear punch injuries at 1-, 2-, 3-, and 4-weeks post-injury. Full wound closure and depigmentation is demonstrated in *Acomys* samples. (B-H) Line plots display ECM proteins and cell markers over the course of wound repair, represented as relative change from baseline (*Acomys,* solid line; *Mus*, dashed line). ECM markers include FGG (B), TNC (C), and COL1 (D). Cell markers include CD36 (platelet, E), CD14 (monocyte, F), CD68 (monocyte/macrophages, G), and CD206 (M2 macrophages, H). Numbers in parentheses represent fold change over baseline at the corresponding time point. Images in Fig 8A are reproduced with permission from Brewer et al^36^.

Analysis of cell marker expression also reveals differing dynamics of cellular infiltration between *Mus* and *Acomys* during wound repair. CD36, a platelet marker, was induced in both *Acomys* and *Mus* following injury (Figure 7E). However, the platelet population remained near baseline levels through week 4 in *Acomys*, while a drastic drop in platelet marker signal was observed at week 3 in *Mus*. This differential response suggests that *Acomys* may maintain a more sustained platelet-mediated wound healing response, which could be beneficial for efficient clot formation and tissue repair. Alternatively, rapid and sustained angiogenesis in the newly regenerated tissue in *Acomys* may capture circulating platelets at the time points sampled. CD14, a monocyte marker, displayed a remarkable difference in induction between the two species (Figure 7F). *Acomys* exhibited over a 50-fold induction over baseline at week 1, which subsequently dropped back to baseline levels at week 2. In contrast, *Mus* showed little to no induction over baseline at any time point (Figure 7F). Additionally, CD68, a pan-macrophage marker, exhibited a 15-fold increase above baseline at week 1 in *Acomys*, which remained elevated throughout the 4-week course (Figure 7G). In contrast, *Mus* displayed a drop in expression below baseline at week 1, followed by a substantial increase to 27-fold above baseline at week 4 (Figure 7G). These findings suggest that *Acomys* experience greater initial macrophage infiltration at week 1, while *Mus* exhibit delayed but persistent macrophage induction, potentially reflecting the persistence of an inflammatory environment in *Mus* during later stages of wound repair. Markers of monocyte/macrophage infiltration indicate a greater and more rapid influx of monocytes and macrophages into the wound site after injury in *Acomys* compared to *Mus*, likely contributing to their enhanced regenerative capacity.

While cell markers indicate greater macrophage contribution to early wound repair in *Acomys,* all macrophages are not created equal in the context of wound healing. M1 macrophages are primarily associated with the early inflammatory response, producing pro-inflammatory cytokines and helping to clear pathogens and debris^100^. In contrast, M2 macrophages are pro-regenerative, secreting anti-inflammatory cytokines and promoting tissue repair by stimulating cell proliferation and angiogenesis^101^. The presence of M2 macrophages, often marked by CD206 expression, is a crucial determinant of effective wound healing and tissue regeneration^102^. In our study, CD206 exhibited significant induction in *Acomys*, with a 3-fold increase over baseline at week 1 which persisted until week 4 (Figure 7H). In stark contrast, no induction of CD206 was observed at any time point in *Mus*, implying greater and more persistent infiltration of pro-regenerative M2 macrophages after injury in *Acomys* (Figure 7H). The sustained presence of M2 macrophages in *Acomys* may play a pivotal role in tissue repair, dampening inflammation, and promoting regenerative processes throughout the wound healing continuum^101,103^. Additionally, the long pentraxin PTX3, a pattern recognition molecule and essential component of humoral innate immunity, has been shown to play a critical role in the orchestration of tissue repair, with PTX3 deficiencies resulting in increased fibrin and collagen deposition during wound repair^104^. In addition to its primary immune function, PTX3 regulates wound healing through a variety of mechanisms^105^, including interaction with plasminogen to drive remodeling of the fibrin-rich inflammatory matrix^106^ and resolution of inflammation via macrophage modulation^107^. The presence of PTX3 was shown to cause reductions in a variety of pro-inflammatory macrophage markers, including IL-1β, TNF-α, and CD86, demonstrating the ability of this protein to repress pro-inflammatory M1 macrophage differentiation^107^. In our data, we observe strong induction of PTX3 at week 4 in *Acomys,* while all other sample groups display low or undetectable abundance (data not shown). Elevated levels of PTX3 in *Acomys* at later time points may aid in the resolution of macrophage-induced inflammation and promote pro-regenerative immune cell activity. These findings provide valuable insights into the biological mechanisms driving the regenerative capacity of *Acomys*, with implications for understanding wound repair and regeneration in other species.

To identify correlations between immune cell infiltration and the resulting composition of ECM in newly generated tissue, we performed correlation analysis and generated CIRCOS plots to represent correlation ratios between *Mus* and *Acomys*. Proteins more correlated with specific ECM signatures in *Acomys* than in *Mus* include CD163, a marker of M2 macrophages which displays much stronger correlations to a variety of ECM remodeling proteases and protease inhibitors, as well as coagulation protein FGB, in *Acomys* (Figure 8A). These correlations reflect earlier infiltration of M2 macrophages during wound healing in *Acomys*, concurrent with coagulation and resulting in more active ECM remodeling.

**Figure 8.**
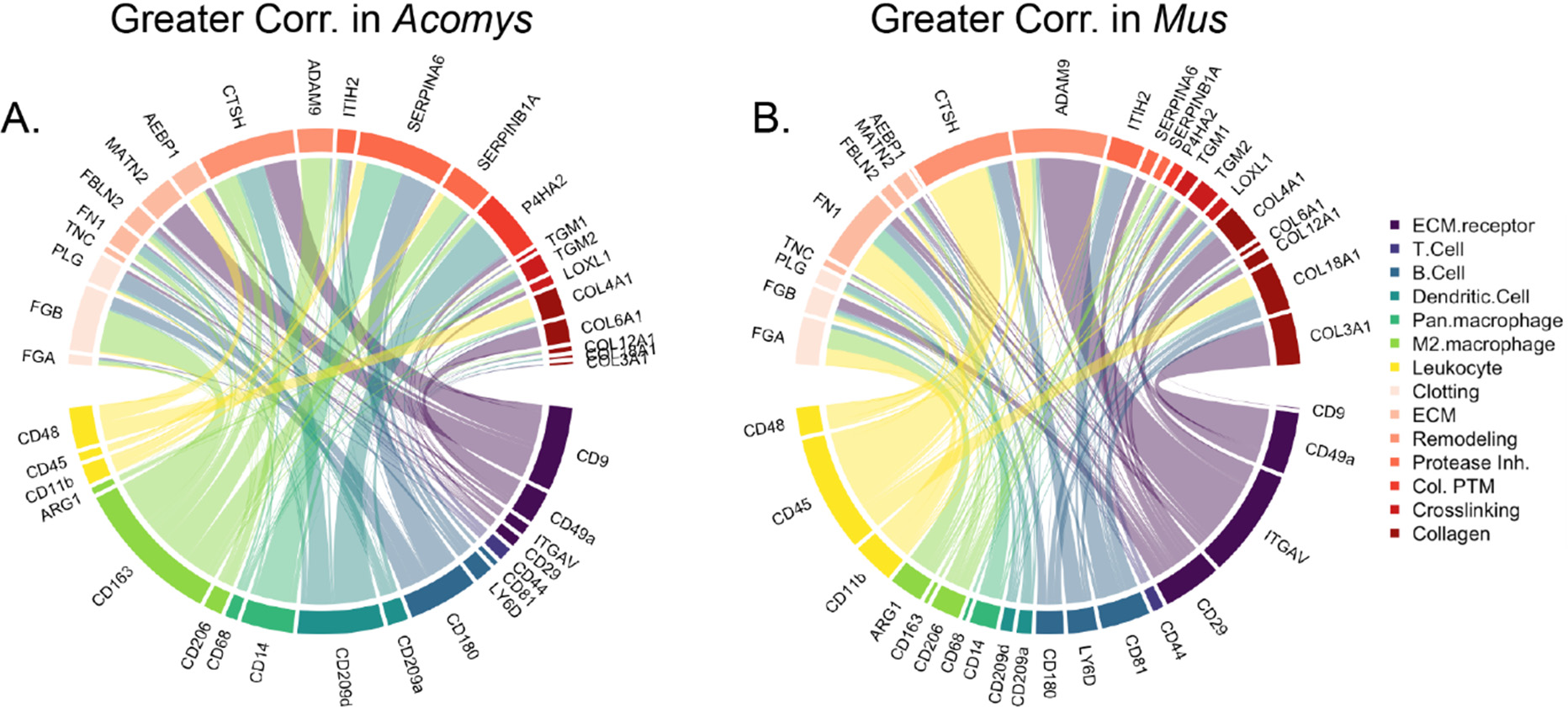
CIRCOS plot displaying correlation ratios between selected cell markers and matrisome components. A) CIRCOS plot displaying *Acomys*/*Mus* correlation ratios. Larger lines indicate protein pairs which are more correlated throughout wound healing in *Acomys* than in *Mus*. B) CIRCOS plot displaying *Mus/Acomys* correlation ratios. Larger lines indicate protein pairs which are more correlated throughout wound healing in *Mus* than in *Acomys*.

In contrast, many protein pairs were additionally observed with greater correlation in *Mus* than in *Acomys* (Figure 8B). Protein pairs with greater correlation in *Mus* include CD45 and CTSH. The expression of CD45, a marker of all leukocytes except mature erythrocytes, strongly correlates with the presence of CTSH, a protease involved in ECM remodeling. While immune cell infiltration is correlated with the presence of ECM remodeling-related proteins in both *Mus* and *Acomys*, the specific correlated immune cell markers and ECM remodelers both vary between the two organisms, suggesting that the process of remodeling is differentially regulated with respect to both timing and molecular activity.

The study of regenerative processes in mammals has tremendous potential to generate insights which result in improved clinical wound repair. In addition to *Acomys*, other models have also contributed to our understanding of scar-free wound healing, including the axolotl (*Ambystoma mexicanum*)^108^, fetal mammals^109–112^, the MRL/MpJ mouse^113^, rabbit ear skin^114,115^, and reindeer antler^116,117^, among others. However, limited proteomics work has been done on these models and most previous studies have inadequately represented ECM proteins. By comparing protein expression patterns identified during *Acomys* regeneration to those observed in other regenerative models, we can potentially identify tissue components which drive regeneration across a variety of mammalian contexts.

The capacity for scar-free regeneration of adult wounds is generally much less common in mammals than it is in other vertebrates^118,119^, appearing in fish^120,121^, amphibians^122^, and salamanders^123^ which have been extensively studied over the past 50 years^124^. One such non-mammalian vertebrate, the axolotl, is able to fully regenerate entire limbs as well as a variety of internal organs, displaying capacity for regeneration which is comparable to that of *Acomys* and greater than that of either frogs or zebrafish^108^. The incredible regenerative ability of both *Acomys* and axolotls provides a unique opportunity for identification of ECM components which contribute to their shared regenerative capacity. A recent review compared expression of ECM-related genes between *Acomys* and axolotl and identified a number of shared expression patterns that may contribute to regeneration in these models, including shared upregulation of COL12A1, TNC, FN1, THBS1, THBS2, and C1QC between the two species^125^. All proteins reported as upregulated in both *Acomys* and axolotl display concurrent upregulation in our data, demonstrating strong agreement between our study and previous analyses. However, analyses incorporated into this review were limited to transcriptomics and analysis of the soluble ECM fraction^125^, limiting the conclusions that can be drawn regarding additional alterations to the insoluble, fibrillar ECM that we are able to capture in this study.

Reindeer antlers are an intriguing model of regeneration due to their rapid growth and extensive remodeling, representing the only mammalian organ that completely regenerates on an annual basis^126^. Independent analysis of both the *Acomys* and reindeer antler models reveals certain protein expression patterns which are shared between the two models during regeneration. COL12A1 is elevated in the regenerating regions of both models, while fibrillar collagens COL1A2 and COL3A1 are reduced both in antler growth centers and in *Acomys* compared to *Mus* after injury^116^. COL12A1 has also been detected at high levels in zebrafish during heart^127^ and spinal cord^128^ regeneration, suggesting a role for this protein in regeneration across species and tissues. In addition to observed shared expression patterns, reindeer antler and *Acomys* also exhibited differences in ECM composition which may indicate variation in regenerative mechanism between these two models. For example, thrombospondin 4 (THBS4) was found to be elevated in the reindeer antler growth center but reduced in *Acomys* healing tissue compared to *Mus*. THBS4 is involved in promoting fibroblast migration and angiogenesis, displaying upregulation during wound repair in *Mus* and *H. sapiens*^129^.. On the other hand, the expression of lumican (LUM) is reduced during reindeer antler regeneration, but induced after injury in *Acomys*^116^. LUM regulates collagen fibrillogenesis and matrix assembly, as well as cell proliferation and migration during wound healing.

A similar injury model to the one used in this paper has been employed since the 1970’s to study regeneration of full-thickness ear punches in rabbit ears. In this model, full-thickness ear punches up to 1 cm in diameter are repaired by the regeneration of new tissue following blastema formation. Comparison of rabbit ear regeneration with *Acomys* ear regeneration has revealed a potential role for endostatin in regulating angiogenesis to improve wound regeneration. Both systemic^130^ and local^131^ injections of Endostar, a recombinant form of human endostatin-XVIII, have been shown to reduce hypertrophic scar formation in rabbit ears. Interestingly, elevated levels of endostatin-XV, but not endostatin-XIII, were observed in *Acomys* during wound regeneration, independent of the parent protein.

Finally, a comparison of fetal scarless wound healing to ear punch regeneration in African spiny mice has revealed some interesting insights into the role of specific proteins in regeneration. In comparison to adult tissue, earlier and more intense deposition of cellular adhesion molecules, including fibronectin (FN1) and tenascin-C (TNC), has been reported to drive fibroblast recruitment and re-epithelialization in human fetal skin repair. We observe elevated levels of both FN1 and TNC in *Acomys* compared to *Mus* at early wound healing time points. Additionally, decorin (DCN), which is involved in the regulation of collagen fibrillogenesis as well as the binding and storage of growth factors within the ECM, is displayed at low levels in injured fetal skin in contrast to injured adult skin. In our data, DCN was observed to have greater expression in *Mus* compared to *Acomys.* While we observe ∼20% greater signal for integrins overall in *Mus*, some integrins, including ITGAV and ITGA5, are specifically upregulated in *Acomys* at week 1. ITGAV and ITGA5 are regulators of adipocyte differentiation, where elevated expression reduces differentiation and increases maintenance of the stem cell population.

Overall, these comparative studies provide important insights into the complex interplay of ECM proteins in scar-free wound healing and open new avenues for future research. Further investigation is needed to fully understand the mechanisms underlying these differences and to determine how they can be harnessed to improve wound healing in humans.

## Conclusions

The research presented here was undertaken to improve understanding of mammalian regeneration through analysis of differences in ECM composition between *Acomys* and *Mus* throughout wound healing. We identified a wide range of significant differences in expression of both cellular and matrix proteins, demonstrating large-scale differences in tissue composition between the two species at baseline and during wound repair. *Acomys* displayed increased production of many proteins that have previously been associated with accelerated or scarless wound healing in other models, alongside reduced expression of fibrillar collagens and other proteins which contribute to tissue tensile strength. Together, these differences from *Mus* represent potential driving factors for both the weakened skin and regenerative capacity observed in *Acomys.* This fibrillar ECM network in *Acomys* was found to be generally less soluble than that of *Mus,* correlating with differences in expression of crosslinking enzymes and potentially aiding in tissue integrity against the reduced presence of normal matrix contributors to tissue strength. We additionally observed rapid and sustained increases in markers of pro-regenerative M2 macrophages in *Acomys,* which likely support a pro-regenerative immune environment and contribute to *Acomys* regenerative capacity. Differences were also detected in matricryptic signaling peptides, with significant elevation of arresten and endostatin-XV observed in *Acomys* independent from expression of their parent proteins. The function of these matrikines in wound healing is not yet well understood, yet these and other insights gained from this study provide candidate matrix components which can potentially be utilized to improve future regenerative therapies.

This study significantly expands on previous proteomic analyses of *Acomys* wound healing, primarily through the use of an ECM-optimized extraction method to access repair-regulating insoluble proteins and the employment of a species-specific protein database to improve proteome coverage over previous studies. While our *Acomys*-specific protein database enabled improved peptide identification compared to the *Mus* database, limitations still exist. The transcriptome-based approach for database generation resulted in a database lacking some *Acomys* proteins due to incomplete transcript coverage. Database entries were missing or low-quality for some ECM proteins of interest, including aggrecan, COL2A1, filaggrin-2, and elastin, limiting the identification quality for these proteins in *Acomys* samples and subsequently preventing accurate comparisons between *Mus* and *Acomys.* Additionally, the database lacks annotated protein isoforms, which could provide deeper insights into isoform-specific expression changes during regeneration. Future construction of a more comprehensive *Acomys* protein sequence database would aid more complete characterization of the *Acomys* proteome and enhance understanding of its remarkable regenerative abilities.

Although the time points sampled here were chosen to represent as much of the wound repair timeline as feasible, additional sampling to provide greater time resolution during early wound healing and to track wound remodeling past 4 weeks post-injury would likely reveal insights about tissue repair dynamics that were obscured in the analysis presented here. Our data indicates a robust early inflammatory and clotting response in *Acomys*, however profiling at earlier timepoints (e.g. 1-3 days post-injury) could better capture the dynamics of these immediate healing events. Examining earlier timepoints would allow quantification of ECM and immune components during initial clot formation, immune cell influx, and re-epithelialization in *Acomys* versus *Mus*. Additionally, later timepoints past wound closure could reveal key differences in long-term ECM remodeling and maturation. We observed high LOX levels at week 4 in *Acomys* coinciding with reduced ECM solubility, indicating tissue crosslinking may occur later to re-establish mechanical integrity. Analyzing additional late stage timepoints (6 weeks onward) could determine if ECM solubility in regenerated *Acomys* tissue recovers to uninjured levels during tissue maturation. Later timepoints could also assess how immune cell populations resolve inflammation and facilitate transition from regeneration to homeostatic tissue maintenance. Overall, expanding the time course to very early and late wound healing events could provide a more comprehensive trajectory of scarless healing in *Acomys* compared to scarring repair in Mus. Despite these limitations, the thorough characterization of ECM dynamics during regenerative healing in *Acomys* presented this work significantly expands our understanding of the molecular drivers of mammalian regeneration and scarless wound repair. Ultimately, these findings bring us one step closer to identifying key regulators of tissue regeneration which may hold promise for developing new therapeutic strategies to improve wound healing outcomes.

## Acknowledgements

Work was supported by the NIH (Grant Nos. R33CA183685, RM1GM131968, P01HL152961, and R01HL146519) and the University of Colorado Cancer Center Support Grant (P30CA046934). In addition, by the W.M. Keck Foundation, R01- HL121877; T32-HL007312; and the Loie Power Robinson Stem Cell & Regenerative Medicine Fund.

